# Potential global impact of the N501Y mutation on MHC-II presentation and immune escape

**DOI:** 10.1101/2021.02.02.429431

**Authors:** Andrea Castro, Hannah Carter, Maurizio Zanetti

## Abstract

The B.1.1.7 SARS-CoV-2 variant, characterized by the N501Y mutation, is rapidly emerging, raising concerns about its effectiveness on natural as well as vaccine-induced adaptive viral immunity at the population level. Since CD4 T cell responses are of critical importance to the antibody response, we examined the global effects of N501Y mutation on MHC-II presentation compared to the N501 wildtype and found poorer presentation across the majority of MHC-II alleles. This suggests that the N501Y mutation may not only diminish binding of antibodies to the RBD but also interfere with their production by weakening the cooperation between T and B cells, facilitating immune escape.

## Main text

The coronavirus disease 2019 (COVID-19) pandemic is a global public health emergency. Records show 103 million infection cases worldwide with a global death toll of 2.24 million people. Beside lockdown, protective masks and social distancing, the containment of community spread depends on immunity by the individual and the population. SARS-CoV-2 promotes seroconversion by day 7 with IgG antibodies outlasting virus detection but declining within 2-3 months after infection (Long et al. 2020), with little residual immunity after 6 months (Rydyznski Moderbacher et al. 2020). Virus-reactive CD4 T cells are found in ≥70% of COVID-19 convalescent patients (Grifoni et al. 2020; Braun et al. 2020), and show a positive correlation with anti-viral IgG titers. Virus-reactive CD4 T cells are detected in ∼40-60% of unexposed individuals, suggesting cross-reactivity between “common cold” coronaviruses and SARS-CoV-2 (Grifoni et al. 2020; Braun et al. 2020).

On December 19, a new variant of the SARS-CoV-2 virus (B.1.1.7) was reported in the UK (Rambaut et al. 2020). This variant has been detected in over 50 countries and is estimated to be ∼50% more transmissible; the CDC has predicted B.1.1.7 will become the dominant strain by March (Corum and Zimmer 2021; Kupferschmidt 2021a). A key feature of the B.1.1.7 variant is the mutation N501Y (asparagine to tyrosine) in the spike protein’s Receptor Binding Domain (RBD). Critically, this mutation is a key feature in many other variants from Australia, Brazil, Denmark, Japan, the Netherlands, South Africa, Wales, and US states Illinois, Louisiana, Ohio and Texas (Corum and Zimmer 2021). A reason for its rapid spread likely lies on the fact that mutations at residue N501 (e.g. N501F and N501T) enhance binding of SARS-CoV-2 to the ACE2 receptor (Starr et al. 2020) explaining why the N501Y variant may become the dominant strain of the pandemic in the months ahead. Recently, however, the concern has been raised that virus variants may also elude the immune response (Callaway 2021; Kupferschmidt 2021b). How might these mutations contribute to immune escape?

A first obvious possibility is that since residue N501 is embedded in the epitope recognized by numerous neutralizing antibodies isolated in convalescent patients (see (Castro et al. 2020)) antibodies to wildtype RBD may have weaker binding to virus variants, consistent with increased infectivity and virulence in a mouse model. This effect may be amplified by a mutation (E484K) co-expressed with N501Y in the South African and Brazilian variants. However, immune escape could have an alternative explanation.

The adaptive immune response is mediated by B and T cells and is orchestrated as a chain of events with the highly polymorphic Major Histocompatibility Complex (MHC) acting as the master regulator. Presentation of antigen peptides complexed with the MHC initiates the expansion of T cells with direct anti-pathogen effector function, and CD4 T cells that help the activation of B cells producing anti-virus antibodies: T-B cooperation. Reasoning that T-B cooperation favors preferential pairing of B and T cell epitopes (Castro et al. 2020), we thought that amino acids surrounding B cell epitope(s) at the RBD:ACE2 interface would also be required to bind the MHC Class II (MHC-II) to enable T-B cooperation to activate CD4 T cells that provide preferential help to B cells producing neutralizing antibodies against the RBD. This could ultimately bias production toward less effective antibodies with more limited neutralizing capacity. What are the implications of the N501Y mutation on MHC-II binding at the global population level?

We applied NetMHCIIpan-v4.0 (Reynisson et al. 2020), which uses eluted ligand mass spectrometry data, to predict peptide-MHC-II presentation affinities for all 15mers spanning the 501 position, taking the best rank out of 15 spanning peptides as a presentation score. We found that N501Y mutant has poorer binding potential for 4786/5620 (85.2%) of all MHC-II alleles and 1580/1911 (82.7%) of common MHC-II alleles relative to N501 (Figure 1A-C). In addition, 21.8% of all alleles and 21.9% of common alleles are no longer able to present the mutated peptides using a generous binding threshold of 20. A simulation of all possible mutations of N501, including N501F and N501T, shows variable effects on MHC-II binding consistent with its polymorphism (Figure 1D-E).

**Figure 1.**
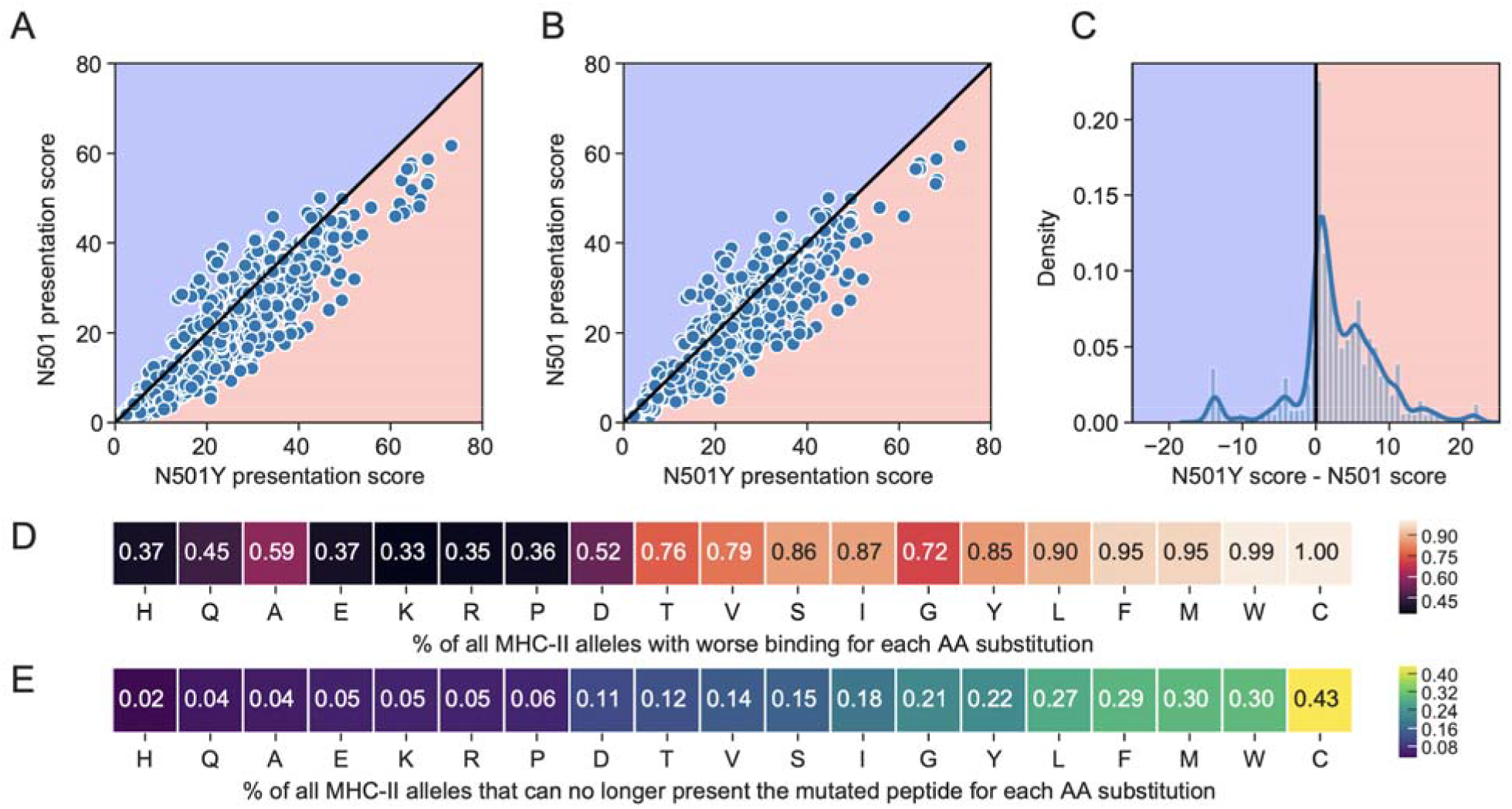
Comparison of percentile rank peptide-MHC binding scores for 15mer peptides spanning the N501 or N501Y position in the spike protein. The blue background indicates improved binding affinity (lower percentile rank) for a N501 peptide versus the corresponding N501Y mutated peptide, and a pink background indicates poorer binding affinity (higher percentile rank) for a N501 vs N501Y peptide. Scatterplots are shown for (A) all available 5620 MHC-II alleles, and (B) the 1911 most common MHC-II alleles. (C) Distribution of N501Y minus N501 MHC-II presentation scores across all MHC-II alleles. (D-E) Heatmaps for all possible single amino acid substitutions at position N501, showing the percent of MHC-II alleles that have (D) worsened binding, or (E) can no longer present the mutated peptide at a threshold of 20.

We provisionally conclude that at the global population level, peptides spanning the N501Y mutation have a generalized lower MHC-II binding affinity for all known and 1911 most frequent alleles. This implies that the N501Y mutation may facilitate the rapid spread of SARS-CoV-2 variants not only by increasing ACE2 binding affinity (Starr et al. 2020) but also by decreasing MHC-II binding for many alleles, weakening preferential T-B cooperation, which is key to producing neutralizing antibodies targeting the RBD. Thus, the new virus variants with the critical N501Y mutation may pose challenges to the quality of the adaptive response required for protection.

This alternative possibility for immune escape to variants carrying the N501Y mutation bears implication for natural immunity to infection and response to current vaccines. Beside proving that antibodies induced by natural infection against the original unmutated virus and by current vaccines recognize and efficiently neutralize virus variants with the N501Y mutation, it will also be important to determine the effect of this mutation on the strength and specificity of neutralizing antibodies in humans. To this, it will also be important to consider the original antigenic sin (Francis 1960) whereby CD4 T cell immunity generated by vaccination against the wild-type spike may be prejudiced to wildtype N501 memory responses favoring immune escape.

## References

Braun, Julian, Lucie Loyal, Marco Frentsch, Daniel Wendisch, Philipp Georg, Florian Kurth, Stefan Hippenstiel, et al. 2020. “SARS-CoV-2-Reactive T Cells in Healthy Donors and Patients with COVID-19.” Nature, July. https://doi.org/10.1038/s41586-020-2598-9.

Callaway, Ewen. 2021. “Fast-Spreading COVID Variant Can Elude Immune Responses.” Nature 589 (7843): 500–501.

Castro, Andrea, Kivilcim Ozturk, Maurizio Zanetti, and Hannah Carter. 2020. “In silico analysis suggests less effective MHC-II presentation of SARS-CoV-2 RBM peptides: implication for neutralizing antibody responses.” PLOS ONE, February. (in press)

Corum, Jonathan, and Carl Zimmer. 2021. “Inside the B.1.1.7 Coronavirus Variant.” The New York Times, January 18, 2021. https://www.nytimes.com/interactive/2021/health/coronavirus-mutations-B117-variant.html.

Francis, Thomas. 1960. “On the Doctrine of Original Antigenic Sin.” Proceedings of the American Philosophical Society 104 (6): 572–78.

Grifoni, Alba, Daniela Weiskopf, Sydney I. Ramirez, Jose Mateus, Jennifer M. Dan, Carolyn Rydyznski Moderbacher, Stephen A. Rawlings, et al. 2020. “Targets of T Cell Responses to SARS-CoV-2 Coronavirus in Humans with COVID-19 Disease and Unexposed Individuals.” Cell 181 (7): 1489–1501.e15.

Kupferschmidt, Kai. 2021a. “Fast-Spreading U.K. Virus Variant Raises Alarms.” Science 371 (6524): 9–10.

Kupferschmidt, Kai. 2021b. “New Mutations Raise Specter of ‘Immune Escape.’” Science 371 (6527): 329– 30.

Long, Quan-Xin, Xiao-Jun Tang, Qiu-Lin Shi, Qin Li, Hai-Jun Deng, Jun Yuan, Jie-Li Hu, et al. 2020. “Clinical and Immunological Assessment of Asymptomatic SARS-CoV-2 Infections.” Nature Medicine 26 (8): 1200–1204.

Rambaut, Andrew, Nick Loman, Oliver Pybus, Wendy Barclay, Jeff Barrett, Alesandro Carabelli, Tom Connor, Tom Peacock, David L. Robertson, and Volz, Erik, COVID-19 Genomics Consortium UK (CoG-UK). 2020. “Preliminary Genomic Characterisation of an Emergent SARS-CoV-2 Lineage in the UK Defined by a Novel Set of Spike Mutations.” December 18, 2020. https://virological.org/t/preliminary-genomic-characterisation-of-an-emergent-sars-cov-2-lineage-in-the-uk-defined-by-a-novel-set-of-spike-mutations/563.

Reynisson, Birkir, Bruno Alvarez, Sinu Paul, Bjoern Peters, and Morten Nielsen. 2020. “NetMHCpan-4.1 and NetMHCIIpan-4.0: Improved Predictions of MHC Antigen Presentation by Concurrent Motif Deconvolution and Integration of MS MHC Eluted Ligand Data.” Nucleic Acids Research. https://academic.oup.com/nar/advance-article-abstract/doi/10.1093/nar/gkaa379/5837056.

Rydyznski Moderbacher Carolyn, Sydney I. Ramirez, Jennifer M. Dan, Alba Grifoni, Kathryn M. Hastie, Daniela Weiskopf, Simon Belanger, et al. 2020. “Antigen-Specific Adaptive Immunity to SARS-CoV-2 in Acute COVID-19 and Associations with Age and Disease Severity.” Cell, September. https://doi.org/10.1016/j.cell.2020.09.038.

Starr, Tyler N., Allison J. Greaney, Sarah K. Hilton, Daniel Ellis, Katharine H. D. Crawford, Adam S. Dingens, Mary Jane Navarro, et al. 2020. “Deep Mutational Scanning of SARS-CoV-2 Receptor Binding Domain Reveals Constraints on Folding and ACE2 Binding.” Cell 182 (5): 1295–1310.e20.

